# Vancomycin impairs macrophage fungal killing by disrupting mitochondrial morphology and function

**DOI:** 10.1101/2024.06.25.600580

**Authors:** Ebrima Bojang, Lozan Sheriff, Man Shun Fu, Chloe Wellings, Ketema Abdissa, Victoria Stavrou, Callum Clark, Andrew D. Southam, Warwick B. Dunn, David Bending, Jose R. Hombrebueno, Ilse Jacobsen, Sarah Dimeloe, Rebecca A. Hall, Rebecca A. Drummond

## Abstract

Vancomycin is a widely prescribed antibiotic used in the treatment of Gram-positive bacterial infections. We recently showed that this antibiotic disrupted protective anti-fungal immune responses via microbiome dysbiosis, enhancing susceptibility to invasive candidiasis. Antibiotics are an independent risk factor for developing this life-threatening fungal infection, but whether microbiota-independent mechanisms also drive this association is not clear. Here, we show that vancomycin directly impairs macrophage responses to *Candida albicans*, the main causative agent of invasive candidiasis. Vancomycin-treated macrophages were less able to kill *C. albicans* despite normal phagocytosis rates and were hyper-inflammatory and more likely to die during infection. We found that vancomycin bound to macrophage mitochondria, leading to depolarisation, reduced respiratory capacity and a hyper-fragmented morphology associated with increased ROS production. Taken together, this work demonstrates direct effects of vancomycin on mammalian immune cells, helping us to understand pro-inflammatory effects of this drug and how it promotes susceptibility to life-threatening fungal infection.

## Introduction

Antibiotics are an independent risk factor for developing mucosal and systemic infections with *Candida* fungi, most commonly *Candida albicans* (Ricotta et al., 2020). Antibiotics deplete commensal bacteria that may provide protection against these fungal infections. For example, *Lactobacillus* species prevent inflammatory growth of *C. albicans* in the vaginal mucosa, and the depletion of these bacteria may promote the development of vaginitis and recurring *C. albicans* infections (Auger and Joly, 1980; Sobel, 2007). In the gut, commensal *C. albicans* populations are relatively low in abundance in most people but can significantly increase following treatment with antibiotics (Rolling et al., 2021; Seelbinder et al., 2023). These ‘fungal blooms’ have been observed in patients prior to development of an invasive *C. albicans* infection (Zhai et al., 2020), indicating that outgrowth of *C. albicans* in the gut is a significant risk factor for life-threatening systemic infection.

We previously published that antibiotics also impair antifungal immune responses in an organ-specific manner, leading to the development of fungal and bacterial co-infection and increased mortality in the context of antibiotic treatment (Drummond et al., 2022). Immunocompetent mice pre-treated with broad-spectrum antibiotics, but particularly vancomycin, had disrupted Th17 responses in the intestines due to the depletion of segmented filamentous bacteria (SFB) that resulted in increased fungal burdens in the GI tract and escape of commensal bacteria to the periphery (Drummond et al., 2022). In that work, antibiotics appeared to be mediating immune system defects indirectly via changes to the microbiota. However, some antibiotics can directly impair mammalian cell function. For example, bactericidal antibiotics including quinolones and β-lactams were shown to significantly impair mitochondrial respiration and cause oxidative stress within mammalian epithelial cells (Kalghatgi et al., 2013). Some antibiotics have been shown to interfere with the phagocytic function of macrophages (Cifarelli et al., 1982; Yang et al., 2017), but whether antibiotics directly impair antifungal functions of these cells in not known. Given the critical role macrophages play in the innate immune response to *C. albicans* infections (Lionakis et al., 2023), we examined how antibiotic treatment affected macrophage uptake and killing of *C. albicans*. We chose to focus on vancomycin since we previously showed that single treatment with this antibiotic enhances mortality following intravenous challenge with *C. albicans* (Drummond et al., 2022). We found that vancomycin impaired *C. albicans* killing by macrophages without affecting phagocytosis, instead causing mitochondrial defects and an enhanced inflammatory phenotype.

## Results

### Vancomycin impairs fungal killing by macrophages

We previously found that vancomycin disrupted control of fungal infection in the GI tract and increased intestinal permeability, leading to escape of commensal bacteria, which was associated with increased mortality (Drummond et al., 2022). In an independent animal facility, we confirmed that vancomycin-treated mice had increased fungal burdens specifically within the GI tract following intravenous challenge with *C. albicans* (**Fig 1A**). Moreover, we observed increased burdens of bacteria in the spleens of vancomycin-treated mice (**Fig 1B**), in line with our previous observations (Drummond et al., 2022). We previously linked this phenotype with antibiotic-driven dysbiosis that led to disruption of lymphocyte responses within the GI tract, including reduced production of GM-CSF and IL-17 (Drummond et al., 2022). These cytokines act on myeloid cells to activate phagocytosis and antifungal killing pathways (Lionakis et al., 2023). However, antibiotics may also directly affect mammalian cell responses in the absence of a microbiota or lymphocytes (Yang et al., 2017). To determine whether vancomycin has a direct effect on myeloid cells, we moved to an *in vitro* system to test the effects of the antibiotic without confounding factors caused by microbiota dysbiosis and changes in lymphocyte functional phenotype. We differentiated mouse bone-marrow derived macrophages in the presence of vancomycin or in antibiotic-free media and examined macrophage maturation and viability. We found that differentiating macrophages in the presence of vancomycin had no effect on the number of macrophages generated or the expression of activation/maturation markers (**Fig 1C, D**). Next, we challenged the macrophages with live *C. albicans* and measured their fungal killing capacity by comparing yeast cell viability after incubation with the macrophages. We found that vancomycin-treated macrophages had significantly impaired fungal killing, both after long-term exposure to a dose previously measured in vancomycin-treated humans (Hu et al., 2022) of 20 µg/mL for 5 days, and to short-term high dose exposure (100 µg/mL for 6 hours) (**Fig 1E**). These data demonstrate that vancomycin impairs macrophage killing of *C. albicans*.

**Figure 1:**
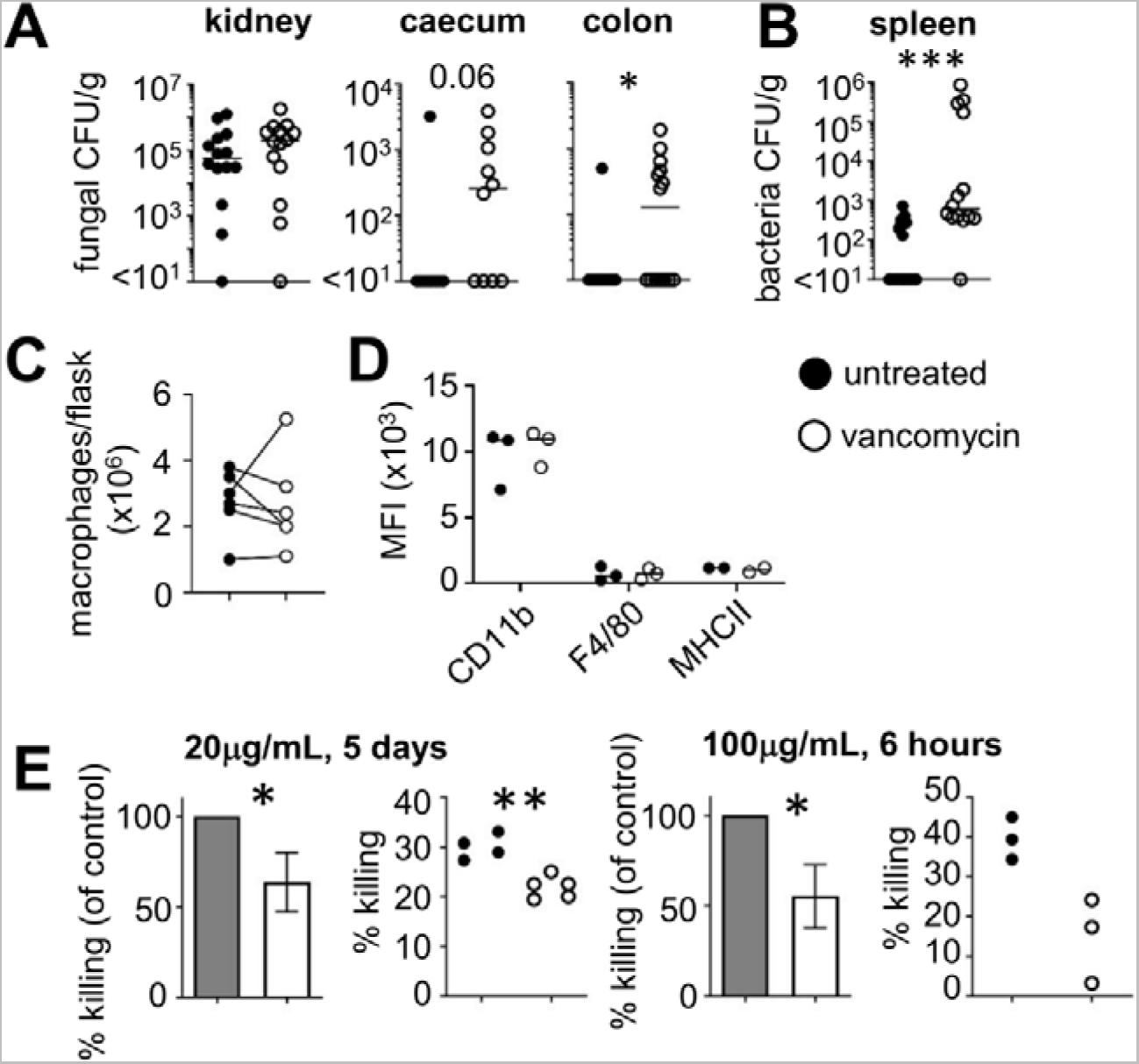
Vancomycin directly impairs fungal killing by macrophages. (A) Wild-type C57BL/6 mice were treated with vancomycin in the drinking water for 4 weeks, or left untreated, prior to intravenous infection with *C. albicans* SC5314. Indicated organs were isolated at day 7 post-infection for fungal or (B) bacterial burden analysis. Each point represents an individual animal. Data pooled from 3 independent experiments and analysed by Mann Whitney U-test. (C) The number of macrophages cultured per flask from mouse bone marrow in the presence or absence of vancomycin. Lines show paired samples that used bone marrow cells from the same animal for macrophage differentiation. (D) Mean fluorescence intensity (MFI) or indicated lineage and activation markers on macrophages cultured without antibiotics (untreated) or 20µg/mL vancomycin during differentiation. Each point represents macrophage cultures generated independently from different animals. (E) Fungal killing ability of untreated and vancomycin-treated macrophages for long-term exposure (left) and short-term exposure (right). Bar graphs represent means from 2-3 pooled independent experiments; dot plots show technical replicates from an example experiment. Data analysed by unpaired t-tests. *P<0.05, **P<0.01, ***P<0.005

### Vancomycin does not impair uptake of live *C. albicans* by macrophages

Antibiotics have been previously shown to disrupt phagocytosis (Cifarelli et al., 1982; Yang et al., 2017), and vancomycin has been described to block autophagy pathways that are involved in uptake of fungi and bacteria (Ha et al., 2014). We therefore explored whether vancomycin disrupted phagocytosis of macrophages, leading to the impaired killing we observed. First, we measured general phagocytic capacity by challenging untreated and vancomycin-treated macrophages with fluorescent particles. This experiment revealed that vancomycin pre-treatment impaired macrophage phagocytosis (**Fig 2A**). To examine potential underlying reasons for this, we measured the abundance of the autophagy protein LC3 in macrophages using western blot. LC3-associated phagocytosis (LAP) is a process involving components of autophagy machinery that enable uptake of microbes and activation of inflammatory responses, including to *C. albicans* since LC3-deficient macrophages are impaired in their ability to kill yeast cells (Nicola et al., 2012). Vancomycin was previously indicated to disrupt LC3 abundance and autophagy pathways (Ha et al., 2014), hence we examined whether similar disruptions occurred in our model. We found that infection of macrophages with *C. albicans* significantly upregulated LC3 abundance, but vancomycin did not alter this across several experiments (**Fig 2B**). Next, we examined whether vancomycin pre-treatment specifically affected *C. albicans* uptake. For that, we performed a flow cytometry-based phagocytosis assay in which macrophages are challenged with fluorescent *C. albicans*, and surface-bound and internalised yeast cells distinguished using a counter-stain with anti-*Candida* antibody. In contrast to our observations with fluorescent particles, we observed no significant difference between untreated and vancomycin-treated macrophages in their uptake or binding of live *C. albicans* (**Fig 2C**). Taken together, these experiments show that vancomycin treatment may generally impair macrophage phagocytosis, but these defects are less relevant for uptake of live *C. albicans,* and reduced phagocytosis does not explain the killing defects of these cells.

**Figure 2:**
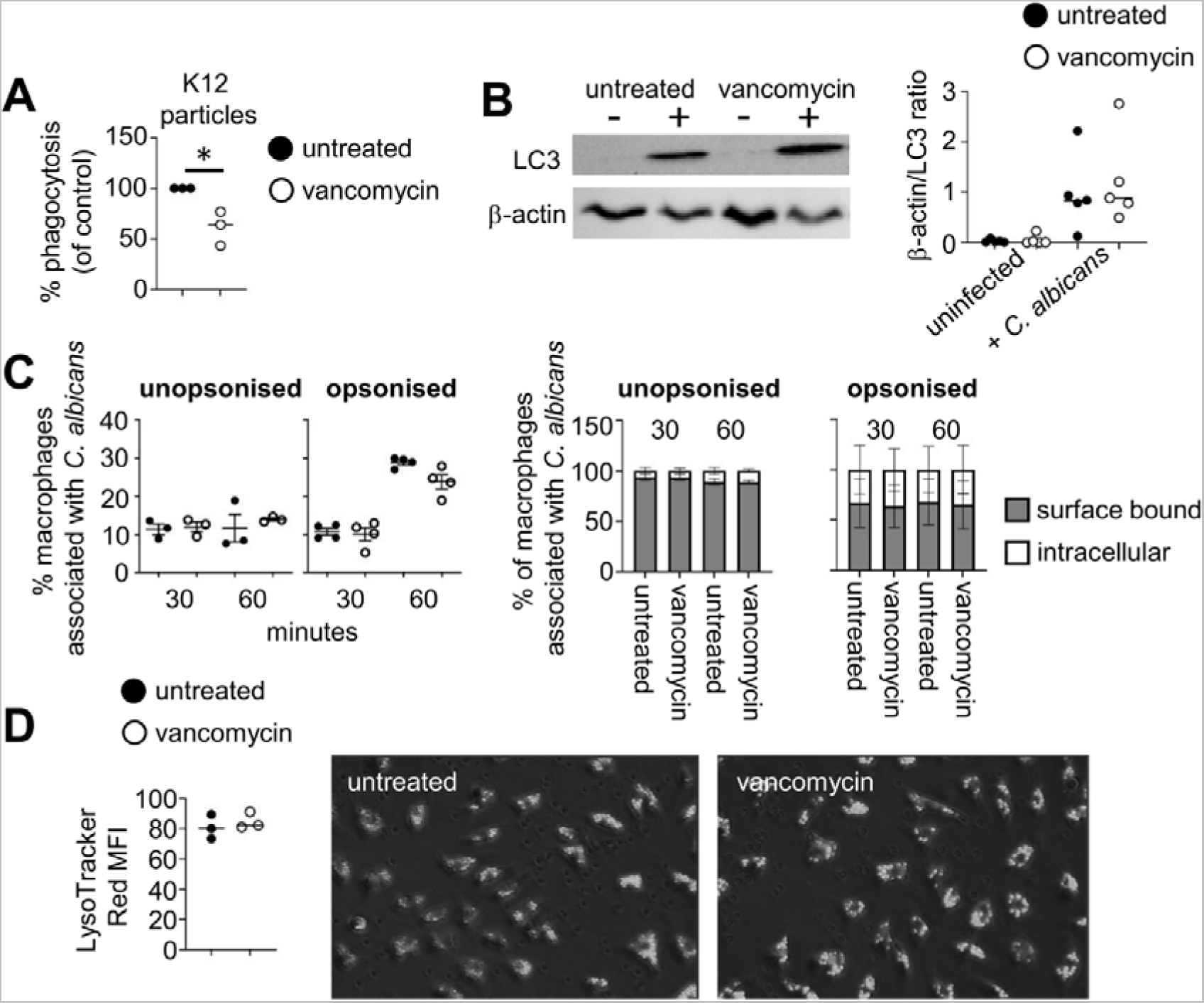
Vancomycin does not affect phagocytosis or phagosome maturation. (A) Phagocytosis assay with K12 *E. coli* particles. Each point represents the mean value of technical replicates from individual experiments using macrophages generated from different animals. Data analysed by unpaired t-test. *P<0.05. (B) Example western blot for LC3 and β-actin from macrophages that were uninfected (-) or infected with live *C. albicans* for 2 hours (+). The graph shows quantification of pixel densities expressed as a ratio between the 2 proteins. Each point represents an individual experiment (n=5 total). (C) Phagocytosis assay (by flow cytometry) with live *C. albicans* that was either unopsonised or opsonised with live mouse serum. Each point represents the mean value of technical replicates from individual experiments using macrophages generated from individual animals. Bar graphs show mean values from individual experiments, shown as mean +/- SEM. (D) Quantification of mean fluorescence of LysoTracker Red staining in macrophages challenge with heat-killed *C. albicans* at 30 minutes post-stimulation. Each point represents an average of 5-6 fields of view from 3 independent experiments. Examples images are from 30 minutes post-stimulation.

### Phagosome maturation is not affected by vancomycin treatment

Following phagocytosis of live *C. albicans*, macrophages must initiate pathways to enable maturation of fungi-containing phagosomes. This includes recruitment of proteins such as Rab GTPases and a reduction in the pH of the phagosomal lumen, which contributes towards pathogen killing and prevents *C. albicans* from forming hyphae that enable escape from the macrophage (Olivier et al., 2022). We examined phagosome maturation in untreated and vancomycin-treated macrophages using the pH-sensitive dye Lysotracker Red to observe the development of acidic environment within fungi-containing phagosomes (**Fig 2D**). These experiments revealed no significant impairment of phagosomal acidification by vancomycin-treated macrophages (**Fig 2D**). Therefore, vancomycin does not impair uptake of *C. albicans* nor prevent the acidification of fungi-containing vacuoles.

### Vancomycin does not directly affect *Candida albicans* growth or hyphae induction

Since vancomycin-treated macrophages had impaired *C. albicans* killing, yet similar phagocytosis rates and phagosome maturation, we considered whether vancomycin could have a direct impact on *C. albicans* and inhibit fungal growth giving rise to what appeared to be a killing defect in our macrophage co-culture assays. To examine this possibility, we incubated vancomycin with *C. albicans* directly and measured the impact on growth, hyphal formation and exposure of chitin in the cell wall. We found that vancomycin had no impact on *C. albicans* growth or on the formation of hyphae (**Fig S1A, B**). Exposure of chitin on the cell wall was also not affected by vancomycin treatment (**Fig S1C**). Therefore, vancomycin does not significantly affect *C. albicans* directly.

### Vancomycin treatment increases expression of inflammatory and oxidative stress genes during fungal infection

Since disrupted phagocytosis and/or phagosome maturation did not explain the killing defect in vancomycin-treated macrophages, we sought to broadly analyse the impact of vancomycin treatment on macrophage responses to *C. albicans* infection using bulk RNA sequencing. We compared untreated and vancomycin-treated macrophages that were either unchallenged or infected with live *C. albicans*. We chose to focus on an early time-point post-infection (2 hours) to circumvent significant cell damage and death caused by *C. albicans* that occurs after this time point (Tucey et al., 2018). Comparison of uninfected control with uninfected vancomycin-treated macrophages revealed that vancomycin-treatment itself did not cause significant transcriptional changes (**Fig 3A** and **Table S1**), in line with our earlier observations that vancomycin treatment did not impair the differentiation or general health of macrophages (**Fig 1**). Upon *C. albicans* infection, we found more substantial transcriptional changes that were shared between untreated and vancomycin-treated macrophages (**Table S2**), including significant upregulation of immediate-early response genes (*Ier3, Junb*), inflammatory immune signalling (*Nfkbia, Pim1, Irf1*), and cytokines and chemokines (*Cxcl2, Tnf*). We next compared the differentially expressed genes (DEGs) between baseline and infection, comparing untreated and vancomycin-treated macrophages. Interestingly, we found that although 76 genes were shared between untreated and vancomycin treated macrophages, there was 44 genes that were uniquely upregulated in untreated control macrophages during infection, and 52 genes that were uniquely upregulated in vancomycin-treated macrophages during infection (**Fig 3B** and **Table S3**). Genes significantly upregulated by control macrophages (but not vancomycin-treated macrophages) included pro-inflammatory surface receptors and transcription factors (*Cd28, Cxcr4, Rora*), glycolytic enzymes (*Eno1, Pfkfb3*), and enzymes involved in biosynthesis of lysosomes and regulation of lipid metabolism (*Tfeb, Olr1, Tmem189*). In contrast, vancomycin-treated macrophages had significant upregulation of genes involved with ubiquitin and oxidative stress responses (*Ubc, Mdm2, Sod2*) and anti-inflammatory genes (*Arg2, Mir155hg, Macir, Nr4a1*). Moreover, we found that vancomycin-treated macrophages had increased expression of *Il1b* and *Nlrp3*, both components of the canonical inflammasome pathway. These results suggested that although vancomycin-treated macrophages have a similar transcriptional response to control macrophages, they have increased expression of inflammatory pathways associated with the inflammasome and concomitant expression of genes involved with dampening inflammation and oxidative stress following infection with *C. albicans*.

**Figure 3:**
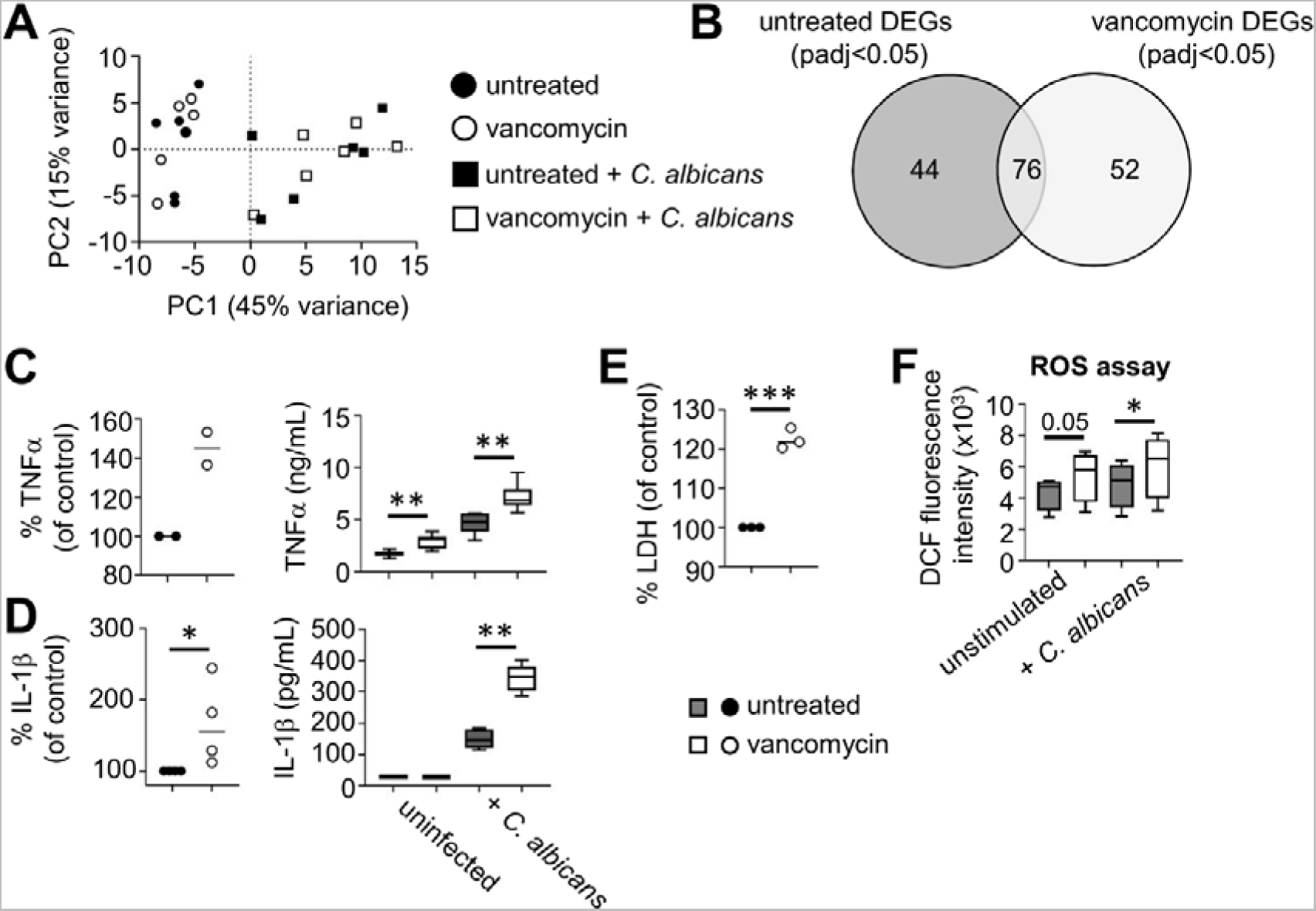
Vancomycin-treated macrophages are more pro-inflammatory. (A) PCA plot of untreated and vancomycin-treated macrophages that were either untreated (circles) or infected with live *C. albicans* for 2 hours (squares) prior to bulk RNA sequencing analysis. Each point represents an individual experiment using macrophages generated from individual animals (n=6 total). (B) Venn diagram comparing the number of unique (44 untreated, 52 vancomycin) and shared (76 genes) differentially-expressed genes (DEGs) between baseline and infection for untreated macrophages (left) and vancomycin-treated macrophages (right). (C) TNFα and (D) IL-1β production by untreated and vancomycin-treated macrophages following infection with live *C. albicans*. The dot plot graphs show mean values for individual experiments, expressed as percentage relative to infected untreated macrophages. The box-and-whisker graphs on the right show technical replicates from an example experiment, showing baseline and post-infection cytokine production. Data analysed by unpaired t-test (pooled data) or two-way ANOVA (example data). (E) LDH assay in LPS-primed macrophages at 4 hours post-infection with *C. albicans*. Each point represents the mean value of technical replicates from individual experiments using macrophages generated from different animals. (F) ROS production assay using H2DCFDA, at baseline and at 3 hours post-infection with heat-killed *C. albicans*. Data is pooled from three independent experiments and analysed by two-way ANOVA. *P<0.05, **P<0.01, ***P<0.005.

To validate our sequencing data, we first analysed production of inflammatory cytokines TNFα and IL-1β to assess the inflammatory status of macrophages and activation of the inflammasome pathway, respectively. In line with our sequencing data, we found a significantly increased production of TNFα by vancomycin-treated macrophages at baseline and following *C. albicans* infection (**Fig 3C**). We found a similar increase in IL-1β production by primed vancomycin-treated macrophages (**Fig 3D**), and an accompanying increase in macrophage death in these conditions (**Fig 3E**). As we had noted an increase in transcripts encoding oxidative stress response genes in vancomycin-treated macrophages, we also examined production of ROS in our macrophages. We found a significant increase in ROS production in vancomycin-treated macrophages following infection with *C. albicans* (**Fig 3F**). Taken together, these data show that vancomycin-treated macrophages have increased production of ROS and inflammatory cytokines following *C. albicans* infection, which may lead to an accelerated death rate under stress.

### Vancomycin causes hyper-fragmentation of mitochondria

Macrophage activation and antimicrobial responses are intimately linked with mitochondrial function (Mills and O’Neill, 2016), and antibiotics are thought to exert direct effects on eukaryotic cells via the mitochondria (Kalghatgi et al., 2013). Mitochondria dysfunction has also been linked with inappropriate inflammation and oxidative stress in epithelial cells (Kalghatgi et al., 2013). Since we observed upregulated expression of genes involved with the regulation of inflammation (including mitochondrial-anchored enzymes *Arg2* and *Mir155hg*) and enhanced ROS production in vancomycin-treated macrophages following *C. albicans* infection (**Fig 3**), we next analysed how mitochondria are affected by vancomycin treatment in macrophages. First, we used confocal microscopy to assess the morphology of mitochondria. After vancomycin pre-treatment, we observed a significant fragmentation of mitochondria and loss of tubular morphology (**Fig 4A**). Mitochondria from vancomycin-treated macrophages were smaller and more rounded compared to control macrophages (**Fig 4B**). To determine the metabolic consequences of this, we analysed mitochondrial respiration using extracellular flux analysis (“Seahorse assay”) comparing untreated and vancomycin-treated macrophages at baseline or when stimulated with heat-killed *C. albicans* (**Fig 4C**). We found a significant reduction in ATP-linked respiration by vancomycin-treated macrophages, but otherwise baseline respiration appeared similar to untreated macrophages (**Fig 4D**). Upon stimulation with *C. albicans*, we found a significant reduction in the spare respiratory capacity of vancomycin-treated macrophages, and consistently, a trend towards reduced maximal respiration (**Fig 4D**). To further examine mitochondrial function, we measured abundance of metabolites of the citric acid cycle before and after stimulation with heat-killed *C. albicans*. Those experiments showed similar abundance of metabolites between untreated and vancomycin-treated macrophages (**Fig S2**), indicating activity of the citric acid cycle was not impaired by vancomycin treatment, in agreement with unaltered basal respiration. We also found no difference in baseline extracellular acidification rate (ECAR) and lactate secretion in vancomycin-treated macrophages (**Fig 4E,F**), indicating that glycolytic function was unaffected by vancomycin. Taken together, these data show that vancomycin significantly alters mitochondrial morphology in macrophages and their oxidative capacity when challenged with *C. albicans*.

**Figure 4:**
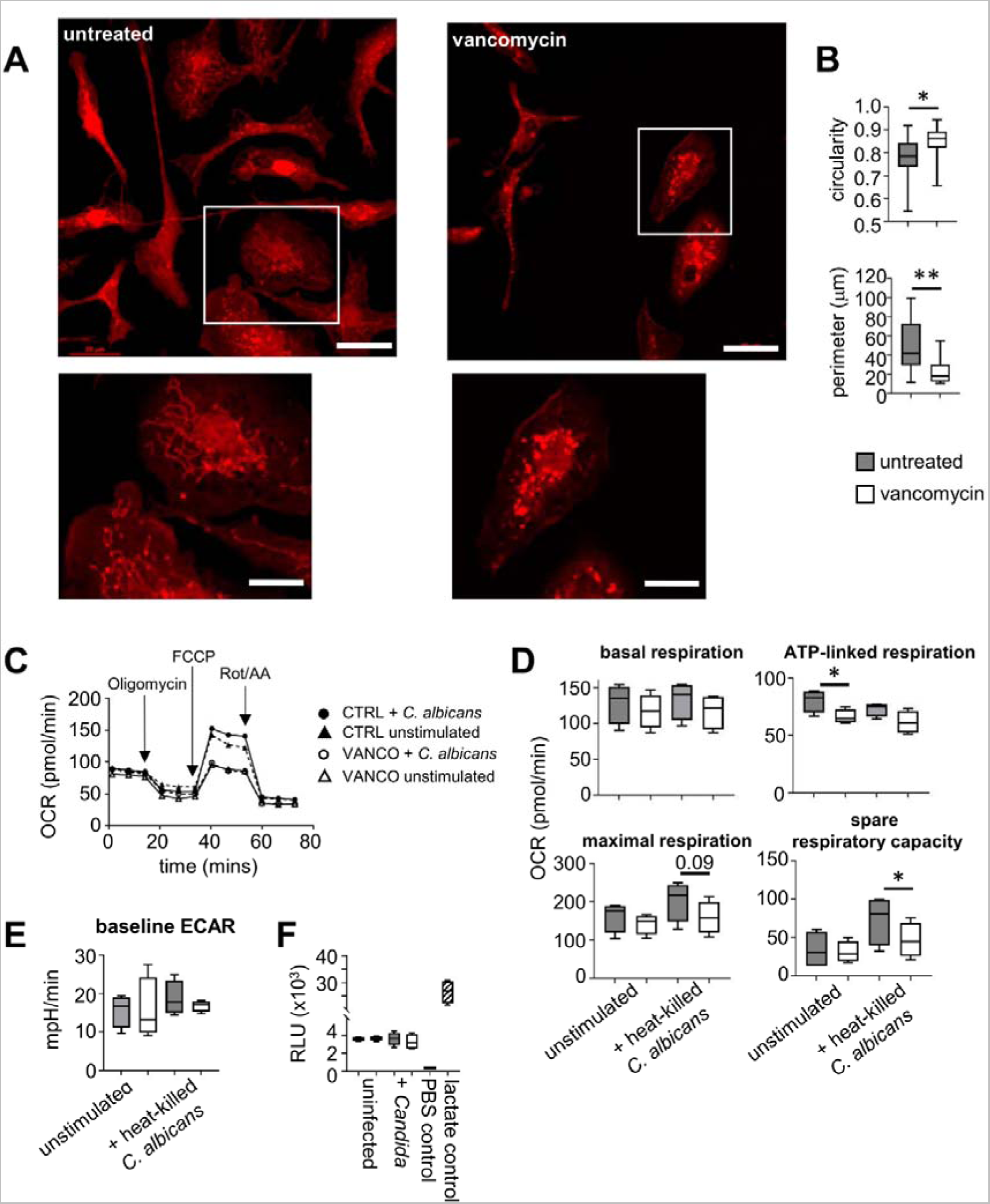
Mitochondria are hyper-fragmented following vancomycin treatment. (A) Untreated and vancomycin-treated macrophages were stained with MitoTracker Red to visualize mitochondria morphology and imaged using confocal microscopy. (B) Images were quantified to determine the circularity of mitochondria and perimeter of mitochondria. Plots show pooled data analysing 200-500 macrophages from 3 individual experiments. (C) Example Seahorse trace from an individual experiment comparing unstimulated macrophages or macrophages stimulated with heat-killed *C. albicans*. (D) Quantification of Seahorse data pooled from 4 individual experiments and analysed by paired t-tests, *P<0.05. (E) Baseline ECAR determined by Seahorse assay. Data pooled from 4 individual experiments. (F) Lactate release assay comparing macrophages at baseline or at 4 hours post-infection with live *C. albicans*. Data is from a single representative experiment (4 technical replicates) that was repeated 2 times.

### Vancomycin penetrates macrophages and localises to mitochondria

We next sought to understand how vancomycin might be exerting its effects on mitochondria by examining its localisation within cells. For this, we exposed macrophages to fluorescently labelled vancomycin and used confocal microscopy to determine the cellular localisation of vancomycin binding. We observed that vancomycin strongly co-localised with the mitochondria and nucleus, as well as the plasma membrane (**Fig 5A**). Vancomycin has previously been shown to have an affinity for the lipid phosphatidylglycerol [PG] (Hu and Tam, 2017), a lipid species found within the inner mitochondrial membrane (Schenkel and Bakovic, 2014). PG is the precursor to the mitochondrial-specific lipid cardiolipin, which plays crucial roles in stabilising the electron transport chain and cristae formation within mitochondria (Falabella et al., 2021). We therefore explored how vancomycin exposure might affect abundance of cardiolipin using a fluorescence-based probe assay, but found no effect of either vancomycin treatment or *C. albicans* infection on cardiolipin levels (**Fig 5B**). Taken together, these data show that vancomycin can penetrate macrophages and localise to mitochondria, likely via it’s lipid-binding affinities to PG, but does not significantly alter abundance of cardiolipin, a mitochondrial-specific lipid, within macrophages.

**Figure 5:**
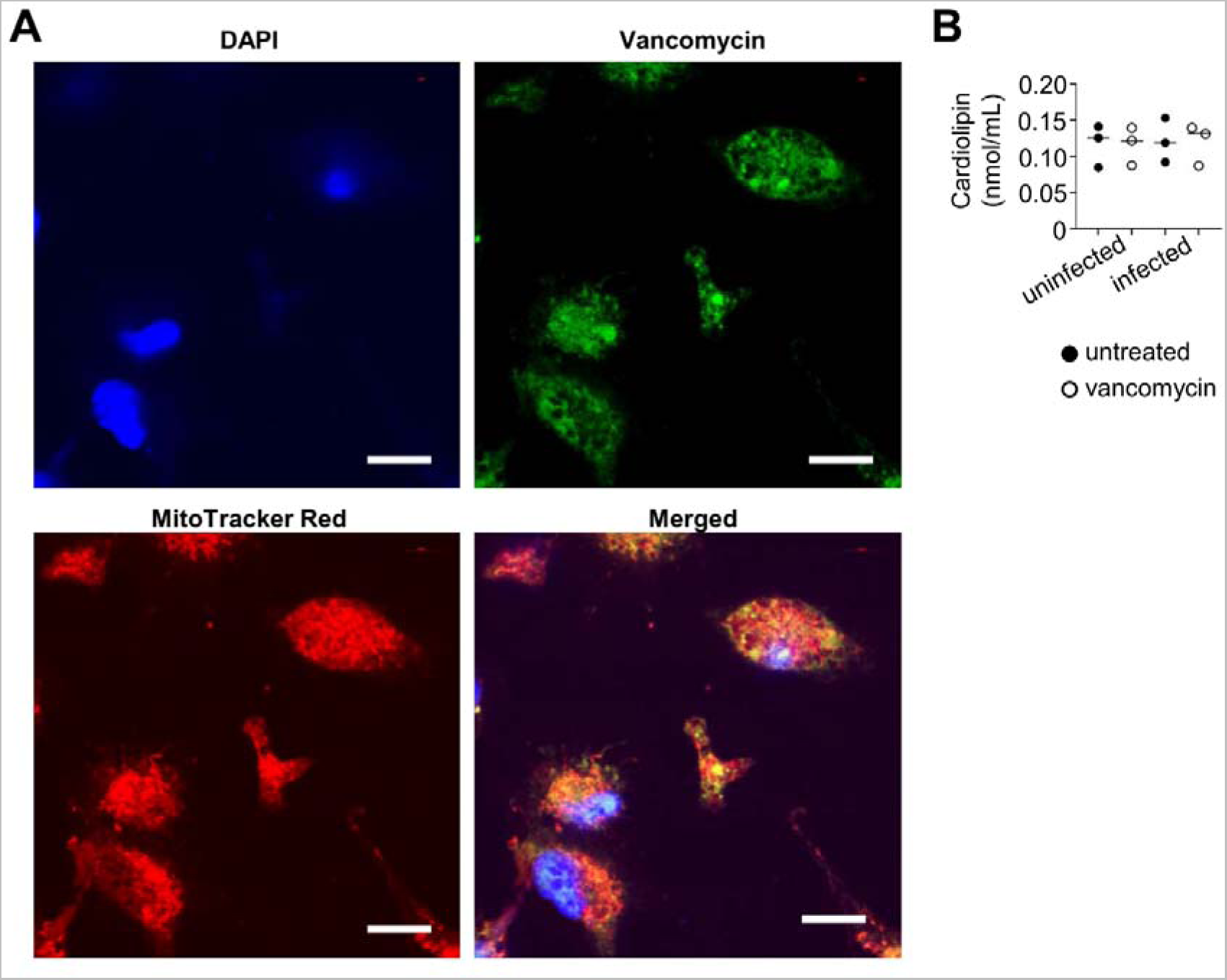
Vancomycin binds to macrophage mitochondria. (A) Example confocal microscopy images of macrophages stained with BODIPY-FL vancomycin (50µg/mL for 4-5 hours), MitoTracker Red and DAPI. Scale bare is 10µm. (B) Cardiolipin abundance in uninfected and infected (MOI 2:1; 2 hours post-infection) untreated or vancomycin-treated macrophages. Each dot represents an individual experiment (n=3 batches of macrophages form individual mice).

### ROS drives mitochondrial depolarisation which is required for fungal killing

Hyper-fragmented mitochondria has previously been associated with oxidative stress and reduced membrane potential and increased ROS generation, linked to prolonged electron dwell time at complexes I and III (Hung et al., 2018; Yu et al., 2006). In line with that, we found increased ROS production by these cells and increased expression of oxidative stress genes (**Fig 3**). To determine if these phenotypes are linked, we first measured the production of mitochondrial-specific ROS in untreated and vancomycin-treated macrophages using mitoSOX staining. These experiments demonstrated that vancomycin treatment induced significantly more mitochondrial ROS both at baseline and during infection (**Fig 6A**). Oxidative stress, caused by over-production of ROS, can occur at both extremes of mitochondrial membrane potential as the cell attempts to balance antioxidant and oxidative redox couples (Li et al., 2013). We therefore examined the impact of vancomycin treatment on mitochondria membrane potential (ψm) using the JC-1 dye, which exhibits a red fluorescence when accumulated within polarised mitochondria and a green fluorescence within non-polarised mitochondria (Reers et al., 1991). Using this approach, we found that vancomycin significantly dissipated ψm (lower red/green ratio) when compared to control macrophages (**Fig 6B, C**). Indeed, vancomycin-treated macrophages had similar ψm to untreated macrophages exposed to the mitochondrial uncoupler BAM15, and to macrophages treated with the ROS inducer menadione which had a significantly decreased ψm (**Fig 6C**). We next directly probed the relationship between mitochondrial depolarisation and fungal killing by macrophages. For that, we pre-treated macrophages with either BAM15 or menadione and then challenged with live *C. albicans* fungi. These experiments showed that macrophages were impaired in fungal killing when treated with either of these drugs, similar to vancomycin-treated macrophages (**Fig 6D**). Taken together, these data show that impaired fungal killing by vancomycin-treated macrophages is linked to the depolarisation of their mitochondria.

**Figure 6:**
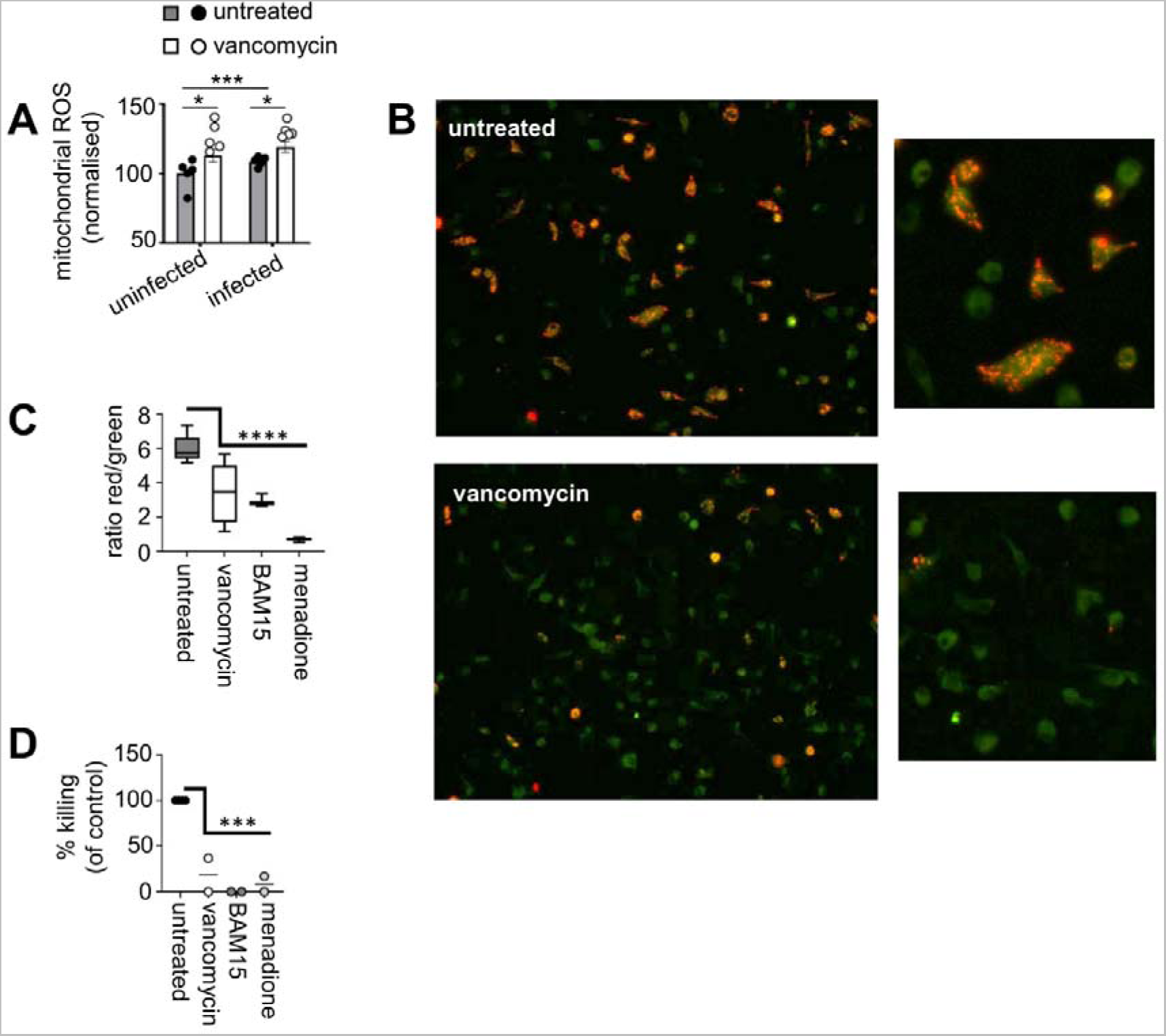
ROS drives mitochondrial polarization which is required for fungal killing. (A) Mitochondrial ROS staining by MitoSOX in untreated and vancomycin-treated macrophages at baseline or at 2 hours post-infection with live *C. albicans*. Bar graphs show mean +/- SEM for 4 independent experiments (using macrophages generated from different animals); overlaid dot plots show technical replicate values for one of these experiments. Data analysed by two-way ANOVA. *P<0.05, ***P<0.005. (B) Example JC-1 labelling of untreated and vancomycin-treated macrophages. (C) Ratio of red and green JC-1 labelling in untreated and vancomycin-treated macrophages, compared to untreated macrophages treated with BAM15 for 2 hours as a positive control and untreated macrophages treated with 5mM menadione for 2 hours. Data pooled from 2 independent experiments in which at least 200 macrophages were analysed. Data analysed by one-way ANOVA, comparing to untreated macrophages. ****P<0.0001. (D) Fungal killing assay using untreated, vancomycin-treated, BAM15-treated and menadione-treated macrophages. Each point represents the mean of technical replicates of individual experiments using macrophages generated from different animals. Data analysed using one-way ANOVA, comparing to untreated macrophages. ***P<0.005.

## Discussion

The immune system is sensitive to antibiotic treatment. Multiple studies have now shown that prior antibiotic exposure affects the ability of the mammalian immune system to fight infections (Deshmukh et al., 2014; Drummond et al., 2022; Erny et al., 2015). Here, we explored the direct effects of vancomycin on anti-fungal immune responses, focusing on macrophages. We found that vancomycin binds to intracellular organelles within mammalian immune cells, which impacted on metabolic capacity and inflammatory function. Vancomycin-treated macrophages had fragmented, depolarised mitochondria and impaired respiratory capacity when exposed to *C. albicans*. These changes to mitochondrial function and oxidative stress associated with decreased capacity of macrophages to kill *C. albicans* fungi.

Vancomycin has been previously shown to disrupt autophagy pathways, resulting in a build-up of LC3 (Ha et al., 2014). LC3-associated phagocytosis (LAP) is a specialised phagocytosis in which components of autophagy pathway are re-purposed for microbial uptake. We found a general phagocytosis defect in vancomycin-treated macrophages when challenged with fluorescent particles made from *E. coli* bacteria. These particles are not live and likely engage LPS receptors such as TLR4. Several antibiotics have been reported to disrupt phagocytosis processes, including ciprofloxacin and ampicillin (Cifarelli et al., 1982; Yang et al., 2017). However, we found that when vancomycin-treated macrophages are challenged with live *C. albicans*, there is no impairment in uptake of yeast compared to untreated control macrophages. Previous work found that LC3 is readily recruited to *C. albicans*-containing phagosomes (Nicola et al., 2012) and LC3-deficient macrophages are impaired in their *C. albicans* killing capacity and were more inflammatory indicated by greater production of TNFα and IL-1β when stimulated with *C. albicans* yeast (Tam et al., 2014). Recruitment of LC3 to *C. albicans*-containing phagosomes depended on recognition of β-glucan via Dectin-1 and ROS generation but did not require TLR signalling (Tam et al., 2014). However, *C. albicans* has also been shown to inhibit LC3 turnover within macrophages and may interfere with deposition of this protein on phagosomes (Duan et al., 2018). Indeed, LC3 recruitment to phagosomes is increased in macrophages that have phagocytosed heat-killed yeast compared to live yeast (Tam et al., 2014). In line with these findings, we observed an increase in LC3 abundance in macrophages infected with *C. albicans*, but this was unaffected by vancomycin-treatment. Therefore, while vancomycin may impair phagocytosis generally this may be less relevant for the fungal pathogen *C. albicans*, which is able to alter the dynamics of LC3 deposition on phagosomes. Whether vancomycin impairs other specific phagocytosis pathways that are independent of LC3, and if this has an impact on uptake of other types of pathogens (e.g. bacteria), will be important to determine the relevance of phagocytic defects caused by vancomycin in susceptibility to other types of infection.

In contrast to phagocytosis, we found a consistent and profound impact on the mitochondria of vancomycin-treated macrophages. Vancomycin co-localised with mitochondria, which had a significantly altered morphology associated with depolarisation and increased ROS production. Mitochondrial respiration and remodelling help determine myeloid cell function. Macrophages responding to LPS undergo a switch from oxidative phosphorylation to glycolysis to generate energy, resulting in a rewiring of the electron transport chain (ETC) and significant changes to mitochondrial function (Mills and O’Neill, 2016). Indeed, the ability to dynamically alter mitochondrial respiration in response to stimuli and cytokines are required to support myeloid functions such as antigen presentation (Kiritsy et al., 2021), tissue repair (Cai et al., 2023) and clearance of bacterial infection (Garaude et al., 2016). Using fluorescently labelled vancomycin as a probe, we found that vancomycin bound to multiple structures within macrophages including mitochondria and the plasma membrane. Previous work that examined vancomycin binding to liposomes showed that the lipid PG had the greatest affinity for vancomycin binding, but that binding capacity was highly dependent on the head groups of the lipids and the curvature of membrane (Hu and Tam, 2017). PG is transported to the inner mitochondrial membrane and acts as a precursor for cardiolipin biosynthesis, a lipid found exclusively within mitochondria. PG is a relatively rare lipid species in mammalian membranes and is more abundant in bacterial membranes (Sim et al., 2023). Vancomycin may therefore localise to mitochondria in mammalian cells via the increased abundance of PG used for cardiolipin synthesis, however we also saw significant labelling of other membrane structures including the plasma membrane and nucleus. The binding of vancomycin to other lipid types in different cell types, particularly macrophages, will be required to determine how this antibiotic penetrates immune cells and the extent to which this affects function of different organelles.

The composition of membranes differs between intracellular organelles (Sim et al., 2023), and these differences may affect vancomycin binding in different cell types, but this may also be influenced by additional factors such as shape, charge and presence of transmembrane proteins. The function and integrity of mitochondria depends on the composition of lipids that make up their membranes. The ratio of lipid species found within mitochondria is similar between different cell types indicating that the types of lipids found in mitochondria are likely integral to their function (Osman et al., 2011). Changes to the biosynthesis of lipids found within mitochondrial membranes show that respiration and function critically depend on a tight homeostasis in these pathways to maintain the function of these organelles. For example, the lipid PC is involved with protein transport across mitochondrial membranes while PE is critical for maintaining polarisation (Schuler et al., 2016). Future work should aim to closely examine the dynamics of lipid species in macrophages during fungal infection and how this might correlate with functional phenotypes. Indeed, other work has shown that macrophages significantly remodel their ‘lipidome’ when activated with specific cytokines and/or inflammatory signals (Zhang et al., 2017), indicating that these biomolecules are sensitive to inflammatory signals.

Indeed, we observed that vancomycin-treated macrophages were more inflammatory than their untreated counterparts, producing more TNFα and IL-1β when exposed to *C. albicans*, resulting in divergent inflammatory phenotypes. Our RNA sequencing analysis an upregulation of multiple anti-inflammatory genes in vancomycin-treated macrophages upon infection that we did not observe in untreated macrophages. These included the enzyme Arginase-2 and nuclear receptor Nr4a1. Arginase-2 is an anti-inflammatory enzyme anchored to the mitochondria and responsible for downregulating IL-1β and HIF1α signalling in response to IL-10 stimulation (Dowling et al., 2021). Arginase-2 is further regulated by microRNA-155 (Dowling et al., 2021), which we also found was significantly increased in vancomycin-treated macrophages upon *C. albicans* infection. Nr4a1 (also known as Nur77) plays an integral role in limiting macrophage inflammatory responses via its role in mitochondrial respiration. Nr4a1-deficient macrophages have greater production of IL-1β and IL-6 that was associated with a reduction in oxidative phosphorylation and a break in the citric acid cycle leading to a build-up of succinate (Koenis et al., 2018).

Although we observed significant effects of vancomycin treatment on inflammatory phenotype and mitochondrial function, we did not observe any significant change in glycolysis. Macrophages switching to glycolysis for energy production is known as the Warburg effect, and has been observed during several inflammatory conditions (O’Neill et al., 2016). Indeed, other work has shown that macrophages switch to glycolysis and upregulate pathways involved in glucose uptake during *C. albicans* infection, with this metabolic switch rapidly increasing from 3 hours of infection and peaking at 6-9 hours post-infection (Tucey et al., 2018). These studies analysed infection times that were much longer than in the current study, in which we focused on the earlier time-point of 2 hours post-infection as we wished to examine the broad effects of vancomycin on macrophage function and initial interactions with *C. albicans* yeast. The upregulation of glucose uptake and dependency on glucose for energy production by *C. albicans*-infected macrophages plays a central role in the death of these cells during infection (Tucey et al., 2018). The fungus was also found to upregulate glucose uptake pathways when exposed to macrophages, leading to a metabolic competition between macrophages and *C. albicans*, resulting in glucose starvation and death of macrophages once *C. albicans* numbers sufficiently outcompeted the macrophages (Tucey et al., 2018). Indeed, glucose starvation in *C. albicans*-infected macrophages was found to activate the NLRP3 inflammasome and IL-1β production and subsequent inflammatory cell death (pyroptosis). Macrophage death following *C. albicans* infection did not require formation of hyphae by *C. albicans*, but was highly strain-dependent (Tucey et al., 2020). Indeed, the fungal toxin Candidalysin, produced by *C. albicans*, is a potent inflammasome activator (Kasper et al., 2018), but its production and release varies widely between *C. albicans* clinical isolates (Li et al., 2022). In our work, we found that vancomycin treatment appeared to accelerate inflammasome activation during *C. albicans* infection resulting in increased IL-1β production and a higher rate of macrophage death. These effects may be occurring independently of metabolic competition for glucose, since we did not observe changes in ECAR or expression of genes involved in glucose uptake between untreated and vancomycin-treated macrophages, although we can’t rule out effects on other types of metabolic competition that may be occurring between macrophages and *C. albicans*. Future studies should aim to examine how competition for other sources of energy between macrophages and *C. albicans* (e.g. lipids) fuels infection-relevant phenotypic changes, and how these may be affected in the context of antibiotic treatment.

In summary, we have found functional defects in immune cells exposed to vancomycin that impaired the ability to fight pathogenic fungus *C. albicans*. Vancomycin enhanced inflammatory pathways in macrophages and directly impaired mitochondrial function, which was required for optimal fungal killing by macrophages.

## Acknowledgements

We would like to thank support staff at the Technology Hub at the University of Birmingham for their support with sorting, flow cytometry and microscopy experiments. This work was funded by the British Infection Association (awarded to RAD), the Wellcome Trust (PhD studentship awarded to EB, grant number 222389/Z/21/Z), the Biotechnology and Biological Sciences Research Council (BBSRC) and University of Leicester funded Midlands Integrative Biosciences Training Partnership (PhD studentship awarded to CW, grant number BB/T00746X/1), Diabetes UK (awarded to JH), and the Deutsche Forschungsgemeinschaft (DFG, German Research Foundation) within the Cluster of Excellence “Balance of the Microverse” under Germany’s Excellence Strategy EXC 2051 (project467 390713860) awarded to KM and IJ.

## Methods

### Mice

8-12 week old C57BL/6JCrl mice (males and females) were housed in individually ventilated cages under specific pathogen free conditions at the Biomedical Services Unit at the University of Birmingham, and had access to standard chow and drinking water *ad libitum*. Mice were housed under 12 hour light/dark cycle at 20-24LC and 45-65% humidity. Animal studies were approved by the Animal Welfare and Ethical Review Board at the University of Birmingham and UK Home Office under Project Licence PBE275C33.

### *C. albicans* growth and *in vitro* infections

*C. albicans* strains used in this study were SC5314 and SC5314-dTomato. Yeast was routinely grown in YPD broth (2% peptone [Fisher Scientific], 2% glucose [Fisher Scientific], and 1% yeast extract [Sigma]) at 30°C for 18 hours at 200rpm. For infections, yeast cells were washed twice in sterile PBS, counted using haemocytometer, and diluted to required concentrations. In general, live yeast were added to cultured macrophages at a ratio of 2:1 (yeast: macrophages) unless otherwise stated. In some experiments, yeast were heat-killed prior to use in experiments. Heat-killing was performed by incubating washed yeast cultures at 70°C for 30 minutes. Heat-killed yeast were used immediately following cooling on ice.

### Mouse *C. albicans* infection model

Mice were infected intravenously with 1x10^5^ yeast cells, prepared as above, via the lateral tail vein. Mice were monitored daily for weight change and development of clinical symptoms (e.g. hunched posture, hypothermia) and were euthanised by cervical dislocation at indicated analysis time-points, or when humane endpoints (e.g. 20% weight loss, hypothermia) had been reached, whichever occurred earlier. For analysis of tissue fungal burdens, animals were euthanized and organs weighed, homogenized in PBS, and serially diluted before plating onto YPD agar supplemented with Penicillin/Streptomycin (Invitrogen). Intestines were first thoroughly washed in PBS prior to homogenisation and plating. Bacterial burdens from spleen were measured as above, except homogenates were plated onto sheep blood agar (Remel). Colonies were counted after incubation at 37**°**C for 24-48 hours.

### Differentiation of bone-marrow macrophages

Bone marrow was flushed from the femurs and tibias of mice (8–12-week-old C57BL/6JCrl males and females) into sterile PBS supplemented with 2mM EDTA. Collected bone marrow was filtered through a 70μm filter and centrifuged (1500rpm 5 minutes at 4□C) to pellet the cells. The pellet was resuspended in Roswell Park Memorial Institute (RPMI) media with GlutaMAX and HEPES, further supplemented with 20% heat-inactivated foetal bovine serum and 40ng/mL M-CSF (Biolegend). Half of the cell suspension was further supplemented with 20μg/mL vancomycin hydrochloride (Fisher BioReagents). The cell suspension (10mL per flask) was incubated in T75 flasks (Corning) for 3 days at 37C at 5% CO2, before removing the media and replacing with 10mL fresh media (supplemented with FBS and M-CSF with/without vancomycin, as above), before continuing the incubation for a further 2 days. On day 5 of the culture, adherent macrophages were collected by harvesting in ice-cold 2mM EDTA/PBS. Briefly, the media was removed and cell layer flooded with ice-cold 2mM EDTA/PBS before incubation on ice for 5 minutes. Cells were gently lifted with a cell scraper (Corning) and collected by centrifugation. Macrophages were counted using a haemocytometer and Trypan blue exclusion. Macrophages were cultured in RPMI supplemented with 10% FBS without antibiotics or with 20μg/mL vancomycin in subsequent assays.

### Phagocytosis assays

For *C. albicans* phagocytosis assay, macrophages were lifted and counted as above prior to seeding in 24-well plates (2x10^5^ macrophages per well) and rested overnight. Prepared dTomato-expressing *C. albicans* was then added to the macrophages (at an MOI of 2:1) the following day. In some experiments, *C. albicans* was opsonised by incubating the yeast cells with 10% mouse serum (Thermo Fisher) on ice for 30 minutes prior to adding to the macrophages. Incubation of the infected macrophages was performed at 37LC and was continued for 30 minutes or 1 hour, before immediately placing plates on ice to stop phagocytosis. Macrophages were then lifted using 2mM EDTA/PBS as above, and then stained with the following fluorophore-conjugated antibodies on ice for 15-30 minutes in the dark: anti-mouse CD45-PerCP Cy5.5 (clone 30-F11), anti-mouse CD11b-APC Cy7 (clone M1/70), anti-mouse F4/80-APC (clone BM8) (all Biolegend) and anti-Candida albicans-FITC (Invitrogen). Samples were then washed in FACS buffer (PBS supplemented with 0.01% sodium azide and 2.5% w/v bovine serum albumin) and acquired using a 5-lazer BD LSR Fortessa equipped with BD FACSDiva software. Data analysis was performed using FlowJo software v.10.9.0 (Treestar). For analysis of general phagocytic function, macrophages were seeded into 96-well flat-bottom plates (1x10^5^ macrophages per well) as above. Phagocytosis was assessed using the Vybrant phagocytosis assay kit (Thermo) following the manufacturers’ instructions, reading the plate fluorescence at 480/520nm excitation/emission using a FlexStation3 plate reader equipped with SoftMax Pro7 software.

### Fungal killing assay by Alamar Blue viability

Macrophages (5x10^4^ per well) were seeded into 96 well plates as above. In some experiments, macrophages were pre-treated with 5mM menadione for 2 hours, or 1:1000 BAM15 for 2 hours. Media was removed the following day from all wells and replaced with 50μL sterile PBS supplemented with 10% mouse serum. Prepared *C. albicans* was then added to each well at an MOI of 2:1 and the cells incubated at 37LC for 2 hours. After, the plates were centrifuged (2000rpm for 4 minutes) and supernatants removed. 100μL of 0.02% Triton-X 100 in distilled water was added to each well and the plate incubated at room temperature for ∼5 minutes. Plates were centrifuged again, the supernatant removed and each well washed twice with 100μL PBS. After washing, 100μL of 1x Alamar Blue metabolic indicator dye (Thermo) was added to each well and the plate incubated at 37LC for 18 hours. Plate fluorescence was read at 560/590nm excitation/emission using a FlexStation3 plate reader equipped with SoftMax Pro7 software. The number of yeast per well was calculated by comparing to a standard curve of yeast-only wells, and percentage killing determined based on the difference between starting number and final number after co-incubation with macrophages.

### Lysotracker red staining and live cell imaging

2x10^5^ macrophages were seeded per well in 24-well plates as above. Media was replaced the following day with fresh media containing 50ng/ml Lysotracker Red DND-99 (ThermoFisher). Cells were infected with live or heat-killed *Candida* at an MOI of 6:1 and fluorescence was read every 5 minutes for 6 hours on a Zeiss Axio Observer microscope at 40x magnification, fitted with a temperature-controlled chamber set at 37°C with 5% CO_2_.

### RNA sequencing

Macrophages were seeded overnight in 24-well plates at 1x10^6^ cells/well, and infected with *C. albicans* for 2 hours (MOI 2:1) or mock infected with PBS. Media was then removed from wells and 1mL of Trizol (Life Technologies) added to each well. Lysed cells were lifted from the wells, transferred to RNAse-free Eppendorfs and stored at -80°C. Samples were thawed at room temperature and 200μL of chloroform was added to each tube, mixed by inverting several times and allowed to settle for 5 minutes. Samples were centrifuged at 12,000g for 15 minutes at 4°C. The top layer containing RNA was transferred to RNAse-free Eppendorf tubes and an equal volume of 70% ethanol was added to each sample. Samples were then added to Qiagen columns (RNeasy kit, Qiagen) to be washed and perform a DNAse digest step (Qiagen). RNA was eluted in RNAse-free water, and then QC-tested using the Qubit Fluorometer TapeStation. RNA libraries were then prepared using the Lexogen QuantSeq 3’ mRNA-Seq kit according to the manufacturers’ instructions. Illumina sequencing was performed on the NextSeq 500 sequencing platform using the v2.5 150 cycles mid output flow cell, which produced 75 bp single reads. Reads where mapped to the *Mus musculus* reference genome. Read counts were quantified using HTSeq-count. Differentially expressed genes between biological conditions were detected using DESeq2 analysis pipeline in R (P_adj_<0.05). Sequencing data has been deposited in the NCBI GEO repository under accession number GSE269051.

### ELISAs

Macrophages were seeded overnight in 96-well tissue culture plates at 1x10^5^ cells/well prior to infection with *C. albicans* at an MOI of 6:1. In some experiments, macrophages were first primed with 50ng/mL LPS (Sigma) for 2 hours prior to infection (IL-1β assay). Infected macrophages were incubated at 37LC for 4 (IL-1β assay) or 6 (TNFα assay) hours, and supernatants collected and frozen at -20LC. Detection of IL-1β and TNFα was performed using DuoSet ELISA kits (RnD Systems) according to the manufacturers’ instructions. Plate absorbance at 450nm was read using a SpectraMax ABS Plus plate reader, equipped with SoftMax Pro7 software.

### LDH assay

Supernatants from macrophage cultures (see figure legends for specific conditions) were collected and stored at -20LC. LDH was quantified in the supernatants using the CyQUANT LDH Cytotoxicity Assay Kit (Invitrogen) following the manufacturers’ instructions. Plate absorbance was read at 490 nm and 680 nm within 2 hours. LDH activity was calculated by subtracting the 680 nm absorbance value (background signal) from the 490 nm absorbance value.

### Lactate measurements

Lactate was measured in macrophage culture supernatants using the Lactate-Glo Assay (Promega) following manufacturers’ instructions. Luminescence was measured using a FlexStation3 plate reader equipped with SoftMax Pro7 software.

### ROS induction and measurements

Intracellular reactive oxygen species was assessed using the H_2_DCFDA probe (ThermoFisher). Macrophages were seeded overnight in 96-well white opaque plates (Costar) at a concentration of 1x10^5^ cells/well. Cells were gently washed with pre-warmed PBS and H_2_DCFDA at a concentration of 40μM was added for 30 minutes at 37LC. BMDMs were then infected with heat-killed *C. albicans* at the indicated multiplicity of infection in PBS supplemented with 10% FBS and incubated at 37LC with 5% CO2 for the indicated times. Fluorescence intensity of DCF was measured a fluorescence spectrophotometer (FlexStation 3 equipped with SoftMax Pro 7 software) at an excitation and emission wavelengths of 485 nm and 525 nm respectively.

### Confocal microscopy

Macrophages were seeded overnight onto glass cover slips (12 mm) placed within 24-well plates (Corning) for fluorescent imaging at 2x10^5^ cells per well. Coverslips were mounted onto glass slides prior to imaging. For JC-1 staining, media was removed from each well and replaced with sterile PBS to wash the macrophages. The PBS was removed and replaced with fresh RPMI media containing JC-1 dye (Thermo) diluted to 1:10,000. Macrophages were incubated with the dye for 30 minutes at 37LC, then washed again in PBS. Stained macrophages were imaged using an inverted epifluorescence microscope (Olympus IX71) coupled to Retiga R6 CCD digital camera (QImaging, Teledyne) and the red/green ratio of JC-1 labelling quantified using ImageJ software (v.1.54, Fiji). For MitoTracker Red staining, macrophages were seeded onto coverslips as above and stained in pre-warmed RPMI media containing 50ng/mL MitoTracker Deep Red dye (Thermo) for 30 minutes at 37LC. Macrophages were then washed and fixed in 4% paraformaldehyde for 30 minutes at room temperature, protected from light. Macrophages were then stained with DAPI (2μg/mL) for 30 minutes at room temperature prior to mounting onto glass slides with ProLong Gold anti-fade mountant (Thermo). Stained macrophages were imaged using the Airy Scan function of a Zeiss LSM 880 confocal microscope (Zeiss) using the X63 water immersion objective.

### Fluorescent vancomycin labelling

For vancomycin localisation studies, macrophages were seeded and washed as above, and then incubated in fresh RPMI media containing 50µM BODIPY-FL vancomycin (Thermo) and incubated at 37LC for 4-5 hours. Macrophages were then stained with MitoTracker Red and DAPI, as above. Labelled cells were imaged using the Airy Scan function of a Zeiss LSM 880 confocal microscope (Zeiss) using the X63 water immersion objective.

### SDS-PAGE and western blotting

Macrophages were seeded overnight in 24-well plates as above. Cells were then infected with *C. albicans* at MOI of 2:1 (yeast : macrophages) and incubated for 4 hours. Macrophages were lysed in RIPA buffer supplemented with protease and phosphatase inhibitor (Sigma) on ice for 10 minutes. Protein concentration of the lysates were measured using the microplate procedure of the Pierce BCA protein assay kit (Thermo) as per the manufacturers’ instructions. Protein lysates were first boiled in Laemmli loading buffer (Sigma) for 5 minutes at 95°C. Equal amounts of protein were loaded onto 10% acrylamide gels and run at 90V for 90-120 minutes. Separated proteins were then transferred from gels to 0.45mm PVDF membranes (Thermo) using iBlot 2-gel transfer device (Life Technologies). Membranes were incubated in blocking buffer (6% non-fat milk in PBS-Tween 20 or 5% BSA in PBS-Tween 20) for 1 hour at room temperature. Membranes were briefly washed in wash buffer (PBS supplemented with 10mM Tris, 100mM sodium chloride and 0.1% Tween20) and then probed with anti-β-actin or anti-LC3 antibody (Cell Signalling Technologies), diluted 1:1000 in blocking buffer, overnight at 4°C on a shaker platform. Membranes were washed with washing buffer using three 5-minute cycles. Membranes were then probed with mouse anti-HRP (clone 7076s) or rabbit anti-HRP (clone 7074s) secondary antibody (Cell Signalling Technologies) for 2 hours at room temperature. This was followed by extensive washing in wash buffer (5 cycles of at least 5 minutes) and a final wash step in distilled water. Clarity Western ECL Substrate (Bio-Rad) was added to the membranes for 1 minute in the dark, prior to imaging with a Gel-Doc imager system (Bio-Rad).

### Extracellular flux (“Seahorse”) assay

Control and vancomycin treated macrophages were seeded overnight at 37°C at 2x10^5^ cells per well (150μL final volume of antibiotic-free or vancomycin media, as above) in a 96-well Seahorse tissue culture plate (Agilent). The cartridge/utility plate (Agilent) was hydrated by immersing probes in 200μL of distilled water and incubated overnight in a non-CO_2_ 37LC incubator with adequate humidity. XF Calibrant solution (Agilent) was also pre-warmed in a non-CO_2_ incubator overnight. Heat-killed *C. albicans* was added to each well the following day, at an MOI of 10:1 (yeast : macrophage) and incubation at 37LC continued for a further 4 hours. Seahorse media was prepared using XF DMEM base media supplemented with 1mM sodium pyruvate, 10mM glucose and 2mM L-glutamine (all Agilent). Samples were analysed on a Seahorse X-96 Flux analyser equipped with Wave software. Prior to analysis, the analyser was initialised using the pre-prepared cartridge plate after removing the water and replacing with pre-warmed calibrant and loading the utility plate with oligomycin, FCCP and rotenone/antimycin A (all Agilent; Seahorse mitochondrial stress test kit) in the relevant injection ports. The plate containing cells/fungi had media removed and replaced with 180mL Seahorse media prior to adding to the analyser. Drugs were injected as per the Stress Test kit program at final concentrations of 1.5μM oligomycin, 2μM FCCP and 0.5μM rotenone/anti-mycinA.

### Metabolite screen by LC-MS

Macrophages were seeded into 24 well plates overnight (as above), and then challenged with heat-killed *C. albicans* for 2 hours (10:1 fungi to macrophage). Macrophage monolayers were washed with 0.9% sodium chloride solution on ice twice prior to quenching and lifting with 0.5mL 70% acetonitrile. Macrophages were snap-frozen on dry-ice and thawed wet-ice 3 times prior to spinning down; supernatants were collected (approx. 0.5mL per sample) and stored at -80LC prior to analysis. All samples were collected within 20 minutes of being removed from the 37LC incubator. For LC-MS, 200μL aliquots were lyophilised. 100μL aliquots were combined to prepare a pooled QC sample from which 200μL aliquots were lyophilised. Samples were analysed in a random order applying UHPLC-MS as described below.

#### Chemicals

Acetonitrile, methanol, isopropanol and water (HPLC grade) were purchased from Fisher Scientific (Loughborough, U.K.). Formic acid and acetic acid (≥98.0% purity) was purchased from VWR International (Lutterworth, U.K.), and ammonium formate and ammonium acetate (≥98.0% purity) was purchased from Sigma-Aldrich (Poole, U.K).

#### Sample analysis

Each biological sample, QC sample and blank sample were analysed applying two complementary UHPLC-MS assays; a HILIC assay in positive and negative ion modes to study water-soluble metabolites. The samples were maintained at 4°C and analysed applying two Ultra Performance Liquid Chromatography-Mass Spectrometry (UPLC-MS) methods using a Vanquish Liquid Chromatography System UPLC+ (Thermo Fisher Scientific, MA, USA) coupled with a heated electrospray Q Exactive Plus mass spectrometer (Thermo Fisher Scientific, MA, USA). Ten pooled QC samples were analysed at the start of the analytical batch to condition the analytical system. Pooled QC samples were then analysed after every 6th biological sample and twice at the end of the analytical batch. The process (extraction) blank samples were analysed as injection 6 and as the last injection of the batch. All analysis order of biological samples were randomised applying the RAND() function in Microsoft Excel.

##### HILIC assay

Polar extracts were analysed on a Accucore-150-Amide-HILIC column (100 x 2.1 mm, 2.6 μm, Thermo Fisher Scientific, MA, USA). For positive ion mode analysis, mobile phase A consisted of 10 mM ammonium formate and 0.1% formic acid in 95% acetonitrile/water and mobile phase B consisted of 10 mM ammonium formate and 0.1% formic acid in 50% acetonitrile/water. For negative ion mode analysis, mobile phase A consisted of 10 mM ammonium acetate and 0.1% acetic acid in 95% acetonitrile/water and mobile phase B consisted of 10 mM ammonium formate and 0.1% acetic acid in 50% acetonitrile/water. For both positive and negative ion mode the flow rate was set for 0.50 mL.min-1 with the following gradient: t=0.0, 1% B; t=2.1, 1% B; t=4.1, 15% B; t=7.1, 50% B; t=10.1, 95% B; t=11.0, 95% B; t=11.5, 1% B; t=15.0, 1% B, all changes were linear with curve = 5. The column temperature was set to 35°C and the injection volume was 2 μL. Data were acquired in positive and negative ionisation modes separately within the mass range of 70 – 1050 m/z at resolution 70,000 (FWHM at m/z 200). Ion source parameters were set as follows: Sheath gas = 55 arbitrary units, Aux gas = 14 arbitrary units, Sweep gas = 4 arbitrary units, Spray Voltage = 3.2kV (positive ion) / 2.7kV (negative ion), Capillary temp. = 380 °C, Aux gas heater temp. = 440°C. Data dependent MS2 in ‘Discovery mode’ was used for the MS/MS spectra acquisition applying a pooled QC sample for each sample type using the following settings: resolution = 17,500 (FWHM at m/z 200); Isolation width = 3.0 m/z; stepped collision energies (stepped CE) = 20, 40, 100 [positive ion mode] / 40, 60, 130 [negative ion mode]. Spectra were acquired in five different mass ranges with each range acquired for a separate QC sample injection (QC samples 6-10): 70 – 210 m/z; 200 – 310 m/z; 300 – 410 m/z; 400 – 510 m/z; 500 – 1050 m/z. A Thermo ExactiveTune 2.8 SP1 build 2806 was used as instrument control software in both cases and data were acquired in profile mode.

#### Raw data processing and univariate analysis

Vendor format raw data files (.RAW) were converted to the mzML file format using ProteoWizard software. Deconvolution was performed by the XCMS R package (version 3.12 running in RVersion 4.0.5). XCMS was operated applying min peak width (6s); max peak width (30s); ppm(14); mzdiff (0.002); bw (0.25); mzwid (0.01); minfrac (0.2).. A data matrix of peak areas for metabolite features (m/z-retention time pairs) vs. samples were constructed for each assay. Data for the first eight QC samples were removed from the dataset prior to further processing and analysis. Each data matrix was filtered as follows: any feature whose median intensity in the biological samples was <20× its median intensity of the extraction blank samples was removed; any feature present in < 70% of the QC samples was removed; features with RSD ≥ 30% across the pooled QC samples (QC9 to last QC sample analysed) were removed. Putative metabolite annotation applying MS1 data was performed by applying the Python package BEAMSpy (https://github.com/computational-metabolomics/beamspy). The parameters applied were maximum retention time = 2 seconds; grouping method = Spearman Rank (Coefficient threshold = 0.5, P<0.05) and mass error +/ - 5ppm. Metabolite annotation based on the Human Metabolome Database, Kegg – human and LIPIDMAPS with a mass tolerance of +/ - 5ppm was applied. Statistical and pathway enrichment analysis applied to the metabolomics datasets was performed in MetaboAnalyst v5.0. For statistical analysis, data were normalized to total sample response and log10 transformed. Statistical analysis applied one-way ANOVA (p<0.05).

### *C. albicans* growth assays, chitin exposure and hyphal growth measurements

In some experiments, *C. albicans* was grown in the presence of vancomycin for 24 hours. *C. albicans* was seeded into 96 well plates at 5000 yeast/well, in the presence or absence of vancomycin (200-100μg/mL). Yeast growth was monitored by serial readings at OD_600_ using a FLUOstar Omega microplate reader. Data was averaged across 3 wells per growth condition. For chitin staining, yeast cells grown in YPD media, or YPD supplemented with 20, 50 or 100µg/mL vancomycin, were washed and stained with 10μg/mL wheat germ agglutinin (WGA; Invitrogen) on ice for 30 minutes, followed by 3.5μg/mL of calcofluor white (CFW; Sigma-Aldrich) and incubating on ice for 10-15 minutes. Labelled yeast cells were analysed using a BD Fortessa flow cytometer, as above. Total (CFW) or exposed (WGA) chitin staining was calculated using the median fluorescence intensity of the cells. For hyphal measurements, length was measured using ImageJ software.

### Statistics

Statistical analyses were performed using GraphPad Prism 9.0 software. Details of individual tests are included in the figure legends. In general, data were tested for normal distribution by Kolmogorov-Smirnov normality test and analyzed accordingly by unpaired two-tailed *t*-test or Mann Whitney *U*-test. In cases where multiple data sets were analyzed, two-way ANOVA was used with Bonferroni correction. In all cases, *P* values <0.05 were considered significant.

