## Supplemental Figures for "Vancomycin impairs macrophage fungal killing by disrupting mitochondrial morphology and function"

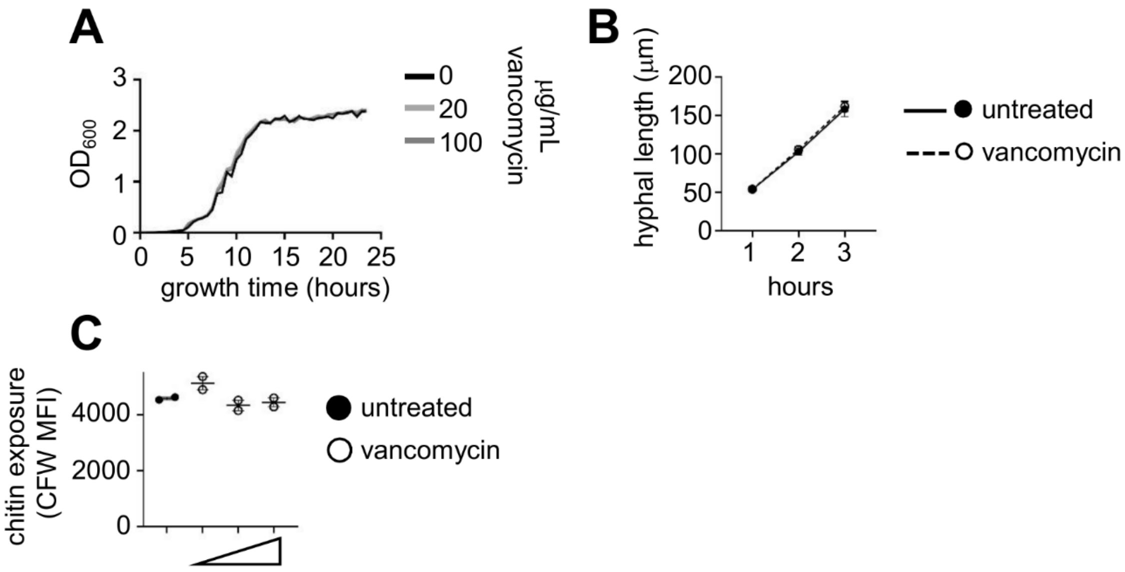


**Figure S1:** (A) *C. albicans* yeast were grown in the presence or absence of vancomycin and growth measured via absorbance at 600nm for 24 hours. Data shown is from a single representative experiment that was repeated 3 times. (B) The average hyphal length at indicated time points following growth in hyphal induction medium. Hyphal lengths were measured from at least 50 cells per time point per condition. (C) Surface chitin exposure as determined by mean fluorescence intensity (MFI) of calcofluor white (CFW) staining and analysis by flow cytometry. Points represent average values from individual experiments. The 3 vancomycin concentrations tested are 20, 50 and 100 µg/mL. Yeast were exposed to vancomycin for 2 hours prior to staining and analysis.

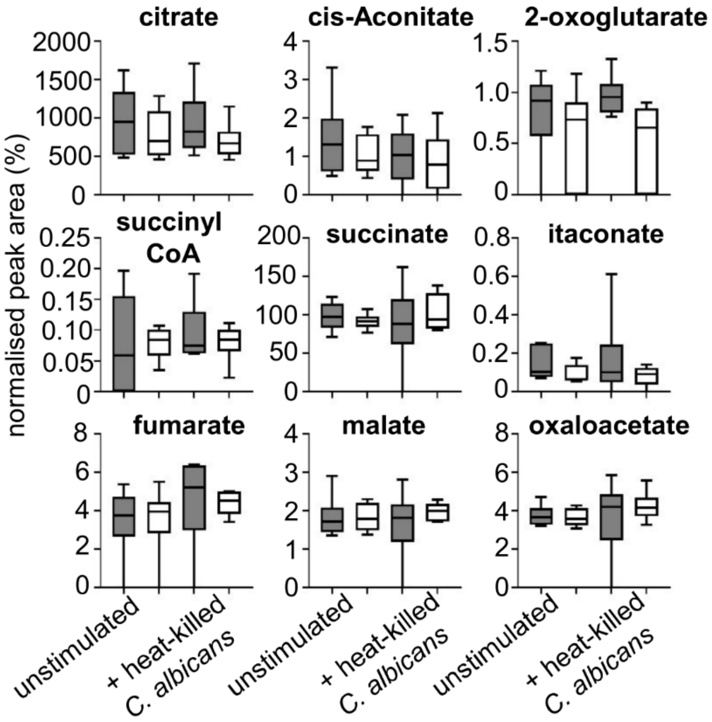


**Figure S2:** LC-MS data measuring levels of indicated citric acid cycle intermediates in untreated (filled bars) or vancomycin-treated (white bars) macrophages unstimulated or treated with heat-killed *C. albicans* for 2 hours. Data pooled from 6 individual experiments (using macrophages generated from individual animals).
